# Metabarcoding lake sediments: taphonomy and representation of contemporary vegetation in environmental DNA (eDNA) records

**DOI:** 10.1101/264903

**Authors:** Inger Greve Alsos, Youri Lammers, Nigel Giles Yoccoz, Tina Jørgensen, Per Sjögren, Ludovic Gielly, Mary E. Edwards

## Abstract

Metabarcoding of lake sediments may reveal current and past biodiversity, but little is known about the degree to which taxa growing in the vegetation are represented in environmental DNA (eDNA) records. We analysed composition of lake and catchment vegetation and vascular plant eDNA at 11 lakes in northern Norway. Out of 489 records of taxa growing within 2 m from the lake shore, 17-49% (mean 31%) of the identifiable taxa recorded were detected with eDNA. Of the 217 eDNA records, 73% and 12% matched taxa recorded in vegetation surveys within 2 m and up to about 50 m away from the lakeshore, respectively, whereas 16% were not recorded in the vegetation surveys of the same lake. The latter include taxa likely overlooked in the vegetation surveys or growing outside the survey area. The percentages detected were 61, 47, 25, and 15 for dominant, common, scattered, and rare taxa, respectively. Similar numbers for aquatic plants were 88, 88, 33 and 62%, respectively. Detection rate and taxonomic resolution varied among plant families and functional groups with good detection of e.g. Ericaceae, Roseaceae, deciduous trees, ferns, club mosses and aquatics. The representation of terrestrial taxa in eDNA depends on both their distance from the sampling site and their abundance and is sufficient for recording vegetation types. For aquatic vegetation, eDNA may be comparable with, or even superior to, in-lake vegetation surveys and therefore be used as an tool for biomonitoring. For reconstruction of terrestrial vegetation, technical improvements and more intensive sampling is needed to detect a higher proportion of rare taxa although DNA of some taxa may never reach the lake sediments due to taphonomical constrains. Nevertheless, eDNA performs similar to conventional methods of pollen and macrofossil analyses and may therefore be an important tool for reconstruction of past vegetation.

## Introduction

Environmental DNA (eDNA), DNA obtained from environmental samples rather than tissue, is a potentially powerful tool in fields such as modern biodiversity assessment, environmental sciences, diet, medicine, archaeology, and paleoecology [1-4]. Its scope has been greatly enlarged by the emergence of metabarcoding: massive parallel next generation DNA sequencing for the simultaneous molecular identification of multiple taxa in a complex sample [5]. The advantages of metabarcoding in estimating species diversity are many. It is cost-effective, it has minimal effect on the environment during sampling, and data production (though not interpretation) is independent of the taxonomic expertise of the investigator [4, 6]. It may even out-perform traditional methods in the detection of individual species [7, 8]. Nevertheless, the discipline is still in its infancy, and we know little about the actual extent to which species diversity is represented in the eDNA records [9, 10]. This study assesses representation of modern vegetation by eDNA from lake sediments.

DNA occurs predominantly within cells but is released to the environment upon cell membrane degradation [4]. It may then bind to sediment components such as refractory organic molecules or grains of quartz, feldspar and clay [11]. It can be detected after river transport over distances of nearly 10 km [9, 12]. When released into the environment, degradation increases exponentially [9, 13], so eDNA from more distant sources is likely to be of low concentrations in a given sample. Once in the environment, preservation ranges from weeks in temperate water, to hundreds of thousands years in dry, frozen sediment [4]. Preservation depends on factors such as temperature, pH, UV-B levels, and thus lake depth [14-16]. Even when present, many factors affect the probability of correct detection of species in environmental samples, for example: the quantity of DNA [8, 17], the DNA extraction and amplification method used [7, 18], PCR and sequencing errors, as well as the reference library and bioinformatics methods applied [4, 18-20]. If preservation conditions are good and the methods applied adequate, most or all species present may be identified and the number of DNA reads may even reflect the biomass of species [6, 7, 21], making this a promising method for biodiversity monitoring.

When applied to late-Quaternary sediments, eDNA analysis may help disclose hitherto inaccessible information, thus providing promising new avenues of palaeoenvironmental reconstruction [22, 23]. Lake sediments are a major source of palaeoenvironmental information [24] and, given good preservation, DNA in lake sediments can provide information on biodiversity change over time [4, 22, 25]. However, sedimentary ancient DNA is still beset by authentication issues [2,10]. For example, the authenticity and source of DNA reported in several recent studies have been questioned [26-30]. As with pollen and macrofossils [31, 32], we need to understand the source of the DNA retrieved from lake sediments and know which portion of the flora is represented in DNA records.

The P6 loop of the plastid DNA *trnL* (UAA) intron [33] is the most widely applied marker for identification of vascular plants in environmental samples such as Pleistocene permafrost samples [34-36], late-Quaternary lake sediments [15, 22, 27, 37-41], sub-modern or modern lake sediments [42], animal faeces [43, 44], and sub-modern or modern soil samples [6, 45]. While some studies include comparator proxies to assess the ability of DNA to represent species diversity (e.g., [35, 41, 46, 47], only one study has explicitly tested how well the floristic composition of eDNA assemblages reflect the composition of extant plant communities [6], and similar tests are urgently needed for lake sediments. Yoccoz *et al.* found most common species and some rare species in the vegetation were represented in the soil eDNA at a subarctic site in northern Norway. The present study attempts a similar vegetation-DNA calibration in relation to lake sediments.

We retrieved sedimentary eDNA and recorded the vegetation at 11 lakes that represent a gradient from boreal to alpine vegetation types in northern Norway. We chose this area because DNA is best preserved in cold environments and because an almost complete reference library is available for the relevant DNA sequences for arctic and boreal taxa [34, 36]. Our aims were to 1) increase our understanding of eDNA taphonomy by determining how abundance in vegetation and distance from lake shore affect the detection of taxa, and 2) examine variation in detection of DNA among lakes and taxa. Based on this, we discuss the potential of eDNA from lake sediments as a proxy for modern and past floristic richness.

## Materials and Methods

### Study sites

Eleven lakes were selected using the following criteria: 1) lakes size within the range of lakes studied for pollen in the region and with limited inflow and outflow streams; 2) a range of vegetation types from boreal forest to alpine heath was represented; and 3) lakes sediments are assumed to be undisturbed by human construction activity. Six of the lakes were selected also for the availability of pollen, macro and/or ancient DNA analyses [27, 48-52]. Data on catchment size, altitude, yearly mean temperature, mean summer temperature and yearly precipitation were gathered using NEVINA (http://nevina.nve.no/) from the Norwegian Water Resources and Energy Directorate (NVE, https://www.nve.no). Lake size was calculated using http://www.norgeibilder.no/. Number and size of inlets and outlets were noted during fieldwork.

### Vegetation surveys

We attempted to record all species growing within 2 m from the lakeshore. This was a practically achievable survey, and data are comparable among sites. Aquatics were surveyed from the boat using a “water binocular” and a long-handled rake, while rowing all around smaller lakes and at least half way around the three largest lakes. We also surveyed a larger part of the catchment vegetation. For this, we used aerial photos (http://www.norgeibilder.no) to identify polygons of relatively homogeneous vegetation (including the area within 2 m). In the field we surveyed each polygon and classified observed species as rare (only a few ramets), scattered (ramets occur throughout but at low abundance), common, or dominant. The area covered and intensity of these broad-scale vegetation surveys varied among lakes due to heterogeneity of the vegetation, catchment size and time constraints. They mainly represent the vegetation within 50 m of the lakeshore. Sites were revisited several times during the growing season to increase the detection rate. For each lake our dataset consisted of a taxon list for 1) the <2-m survey, 2) the extended survey consisting of observations from <2 m and the polygons, 3) an abundance score based on the highest abundance score from any polygon at that lake. Taxonomy follows [53, 54].

### Sampling lake sediments

Surface sediments were collected from the centres of the lakes between September 21^st^ and October 1^st^, 2012, using a Kajak corer (mini gravity corer) modified to hold three core tubes spaced 15 cm apart, each with a diameter of 3 cm and a length of 63 cm (Fig 1, Table 1). The core tubes were washed in Deconex^®^22 LIQ-x and bleached prior to each sampling. The top 8 cm sediments were extruded in field. Samples of ca. 25 mL were taken in 2-cm increments and placed in 50-ml falcon tubes using a sterilized spoon. All samples were frozen until extraction.

**Fig 1.**
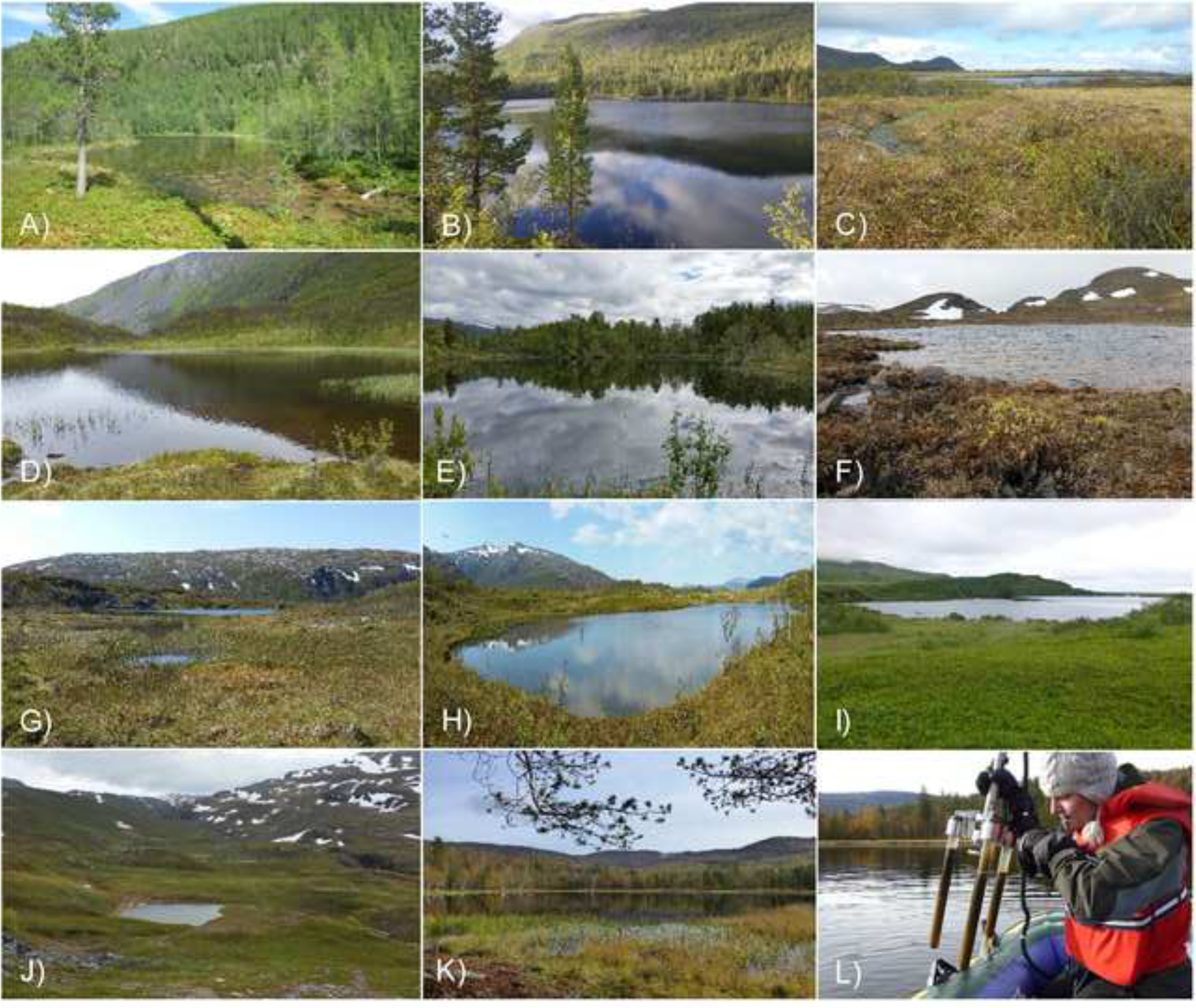
Study lakes in northern Norway. a) A-tjern, b) Brennskogtjørna, c) Einletvatnet, d) Finnvatnet, e) Gauptjern, f) Jula Jävrí, g) Lakselvhøgda, h) Lauvås, i) Øvre Æråsvatnet, j) Paulan Jávri, k) Rottjern, l) Tina Jørgensen sampling surface sediments with Kajak corer. Photo: I.G. Alsos.

**Table 1.** Characteristics of lakes where vegetation surveys and lake sediment DNA analyses were performed.

| Lakes | District | Habitat type | Catchment area (km<sup>2</sup>) | Alt. (m a.s.l.) | Lake size (ha) | Water depth (m) | Yearly mean (°C) | Summer mean (°C) | Yearly prec. (mm) | Inlets | N lat. | E lat. |
| --- | --- | --- | --- | --- | --- | --- | --- | --- | --- | --- | --- | --- |
| A-tjern<sup>a</sup> | Dividalen | Mixed forest/mire, tall herbs | 0.17 | 125 | 1.70 | 5.5 | -0.8 | 6.9 | 636 | 3 | 68.996 | 19.486 |
| Brennskogtjønna | Dividalen | Pine forest, heath | 1.20 | 311 | 10.64 | 20.0 | -0.9 | 6.4 | 457 | 2 | 68.859 | 19.594 |
| Einletvatnet | Andøya | Mires, patches of birch forest | 1.26 | 35 | 27.00 | 4 (6.7) | 3.7 | 8.8 | 1025 | 5 minor | 69.258 | 16.071 |
| Finnvatnet | Kvaløya | Birch forest/mire | 0.20 | 158 | 0.86 | 2.0 | 2.7 | 7.9 | 1005 | 3-4 minor | 69.778 | 18.612 |
| Gauptjern | Dividalen | Sub-alpine mixed forest, tall and low herbs | 0.07 | 400 | 0.79 | 4.0 | -0.9 | 6.5 | 451 | 2 | 68.856 | 19.618 |
| Jula Jávri<sup>c</sup> | Kåfjorddalen | Alpine heath and mire | 1.05 | 791 | 0.04 | 1.7 | -3.6 | 3.9 | 670 | 2-5 minor | 69.365 | 21.099 |
| Lakselvhøgda | Ringvassøya | Alpine heath and mire, scattered birch forest | 0.06 | 143 | 0.77 | 2.0 | 2.5 | 7.2 | 977 | 0 | 69.927 | 18.846 |
| Lauvås | Ringvassøya | Heath, mire and mesic herb birch forest | 0.41 | 4 | 0.71 | 3.3 | 2.7 | 7.5 | 971 | 2 | 69.946 | 18.860 |
| Øvre Æråsvatnet | Andøya | Mires and birch forest, conifers planted | 3.60 | 43 | 24.00 | 9.5 | 3.4 | 8.3 | 1027 | 3 | 69.256 | 16.034 |
| Paulan Jávri | Kåfjorddalen | Alpine heath | 0.56 | 746 | 0.22 | 2.0 | -3.7 | 3.7 | 662 | 1+1 minor | 69.399 | 21.015 |
| Rottjern<sup>b</sup> | Dividalen | Mixed forest, tall herbs | 0.96 | 126 | 1.91 | 3.0 | -0.3 | 7.6 | 619 | 2 | 68.983 | 19.477 |

All lakes are in northern Norway. Water depth given for sampling site in the centre of the lake; deepest point in brackets if different. “Summer” is May-September, “Alt.” is altitude, “prec.” is precipitation, and “N lat.” and “E. lat” are northern and eastern latitude, respectively. Mixed forest is forest dominated by birch but with some Pine.
<sup>a,b</sup>Named A-tjern and B-tjern in Jensen& Vorren 2008. B-tjern later official named Rottjern. A-tjern named “Vesltjønnna” on NEVINA but this name is not official.
<sup>c</sup>Catchment area could not be calculated using NEVINA so this was done in http://norgeskart.no. Temperature and precipitation data were taken from the nearby Goulassaiva.

### DNA extraction and amplification

For each lake, we analysed the top 0-2 cm of sediment separately from two of the three core tubes (n=22). Twenty extra samples from lower in the cores were also analysed. The main down-core results will be presented in a separate paper in which we compare eDNA records with the pollen analyses by [49]. Taxa that were only identified from lower levels in the cores are noted in S1 Table. Samples were thawed in a refrigerator over 24-48 hours, and 4-10 g were subsampled for DNA. The 42 samples and 6 negative controls underwent extraction at the Department for Medical Biology, University of Tromsø, in a room where no previous plant DNA work had been done. A PowerMax Soil DNA Isolation kit (MO BIO Laboratories, Carlsbad, CA, USA) was used following the manufacturer’s instructions, with water bath at 60°C and vortexing for 40 min.

All PCRs were performed at LECA (Laboratoire d’ECologie Alpine, University Grenoble Alpes), using the *g* and *h* universal plant primers for the short and variable P6 loop region of the chloroplast *trnL* (UAA) intron [33]. Primers include a unique flanking sequence of 8 bp at the 5’ end (tag, each primer pair having the same tag) to allow parallel sequencing of multiple samples [55, 56]. PCR and sequencing on an Illumina 2500 HiSeq sequencing platform follows [41], using six PCR negative controls and two positive controls, and six different PCR replicates for each of the 56 samples, giving a total of 336 PCR samples, of which 216 represent the upper 0-2 cm.

### DNA sequences analysis and filtering

Initial filtering steps were done using OBITools [57] following the same criteria as in [41, 42] (S2 Table). We then used *ecotag* program [57] to assign the sequences to taxa by comparing them against a local taxonomic reference library containing 2445 sequences of 815 arctic [34] and 835 boreal [36] vascular plant taxa; the library also contained 455 bryophytes [44]. We also made comparisons with a second reference library generated after running *ecopcr* on the global EMBL database (release r117 from October 2013). Only sequences with 100% match to a reference sequence were kept. We excluded sequences matching bryophytes as we did not include them in the vegetation surveys. We used BLAST (Basic Local Alignment Search Tool) (http://www.ncbi.nlm.nih.gov/blast/) to check for potential wrong assignments of sequences.

When filtering next-generation sequencing data, there is a trade-off between losing true positives (TP, sequences present in the samples and correctly identified) and retaining false positives (FP, sequences that originate from contamination, PCR or sequencing artefacts, or wrong match to database) [17, 20, 58]. We therefore assessed the number of TP and FP when applying different last step filtering criteria. We initially used two spatial levels of comparison with the DNA results: i) data from our vegetation surveys and ii) the regional flora (i.e., species in the county of Nordland and Troms as listed by the Norwegian Bioinformation Centre (http://www.biodiversity.no/). For any lake, both datasets are likely incomplete, as inconspicuous species may be lacking in the regional records [59] and our vegetation surveys did not include the entire catchment area. Nevertheless, the exercise is useful for evaluating how many FPs and TPs are lost by applying different filtering criteria. We defined true positives as sequences that matched a species recorded in the vegetation surveys at the same lake, being aware that this is an under-representation, as the vegetation surveys likely missed species. We defined false positives as species recorded neither in the vegetation surveys nor the regional flora. We tested the effect of different rules of sequence removal: 1) found as ≤1,≤5 or ≤10 reads in a PCR repeat, 2) found as ≤1,≤2 or ≤3 PCR repeats for a lake sample, 3) occurring in more than one of 72 negative control PCR replicates, 4) on average, higher number of PCR repeats in negative controls than in sample, and 5) on average a higher number of reads in negative controls than in samples (S2 Table). The filtering criteria resulting in overall highest number of true positives kept compared to false positives lost were applied to all lakes. These were removing sequences with less than 10 reads, less than 2 PCR repeats in lake samples, and on average a lower number of reads in lake samples than in negative controls.

### Data analyses and statistics

After data filtering, we compared taxon assemblages from DNA amplifications with the taxa recorded in the vegetation surveys. To make this comparison, taxa in the vegetation surveys were lumped according to the taxonomic resolution of the P6 loop (S1 Table), and the comparison was done at the lowest resultant taxonomic level. The majority of results explore only present/absence (taxa richness); quantitative data are given in tables (including Supporting Information).

Multivariate ordinations (Correspondence Analysis and Non-symmetric Correspondence Analysis, the latter giving more weight to abundant species; [60, 61], were run independently on the vegetation data (present/absent using only taxa recorded within 2m) and eDNA data (present/absent). The similarity between ordinations of vegetation and eDNA data was assessed using Procrustes analysis [62], as implemented in the functions procrustes() and protest() in R library vegan [63].

To estimate the percentages of false negatives and positives in the DNA data and in the vegetation survey, we used the approach described in [64]. If we define the probability of a DNA false positive as pDNA_0, the detectability by DNA as p_DNA_1_, the detectability in the vegetation survey as pVEG_1, and the probability that a species is present as pOCC, we can state that the four probabilities of observing Presence(1)/Absence(0) in the DNA and Vegetation are as follows:

1. *Prob(DNA=0,Vegetation=0) = (1-p_OCC_)(1-p_DNA_0_) + p_OCC_ (1-p_DNA_1_) (1-p_VEG_1_)* In this case, if the species is absent in both the DNA and vegetation, it is either absent with probability (1- p_OCC_) and no false positive has occurred with probability (1- p_DNA_0_), or it is present with probability pOCC, but was not detected both in the DNA with probability (1- p_DNA_1_) and in the vegetation with probability (1- p_VEG_1_).
2. *Prob(DNA=0,Vegetation=0) = (1- p_OCC_)(1- p_DNA_0_) + p_OCC_ (1- p_DNA_1_) (1- p_VEG_1_)* In this case, the species is present, not detected in DNA but detected in the vegetation survey.
3. *Prob(DNA=1,Vegetation=0) = (1- pOCC) pDNA_0 + pOCC p_DNA_1_ (1- pVEG_1)* In this case, the species is either absent and is a false DNA positive, or is present, detected by DNA but not in the vegetation survey.
4. *Prob(DNA=1, Vegetation=1) = p_OCC_ p_DNA_1_ p_VEG_1_* In this case, the species is present and is detected both in the DNA and the vegetation survey.

We assumed the four probabilities varied only among lakes, not among species. We also restricted the analyses to species that were detected at least once using DNA, because for species that were never detected using eDNA, different processes might be important. For *p_DNA_1_*, we also considered a model assuming a logistic relationship between *p_DNA_1_* and lake characteristics, such as lake depth or catchment area, that is: logit(*p_DNA_1_*) = b_0_ + b_1_ Lake Covariate. We fitted these models using Bayesian methods, using uninformative priors (uniform distributions on the [0,1] interval) for the false positive/negative rates for DNA, and an informative prior for the detectability in the vegetation survey (uniform prior on the [0.8,1] interval, as detectability was high in the vegetation survey, but we had no repeated surveys or time to detection available to estimate it). We used the R package rjags to run the MCMC simulations [64]. Model convergence was assessed using the Gelman- Rubin statistics [65], values of which were all ∼1.0.

## Results

### Vegetation records

The vegetation surveys provided 2316 observations of 268 taxa, including hybrids, subspecies, and uncertain identifications. Of these, 97 taxa share sequences with one or more other taxa (e.g., 20 taxa of *Carex* and 15 of *Salix).* Another nine taxa were not in the reference library (e.g. *Cicerbita alpina)*, and eight taxa could not be matched due to incomplete identification in the vegetation survey. Eight taxa of *Equisetum* were filtered out due to short sequence length. This left 171 taxa that could potentially be recognized by the technique we used (S1 Table). For the 11 sites, between 31 and 58 taxa were potentially identifiable (Table 2), and this value was positively correlated with vegetation species richness (y=0.67x+10.3, r^2^=0.93, p<0.0001, n=11). Taxonomic resolution at species level was 77-93% (mean 88) and 65-79 (mean 74%) for the <2 m and extended (i.e., combined) vegetation surveys, respectively.

**Table 2.**
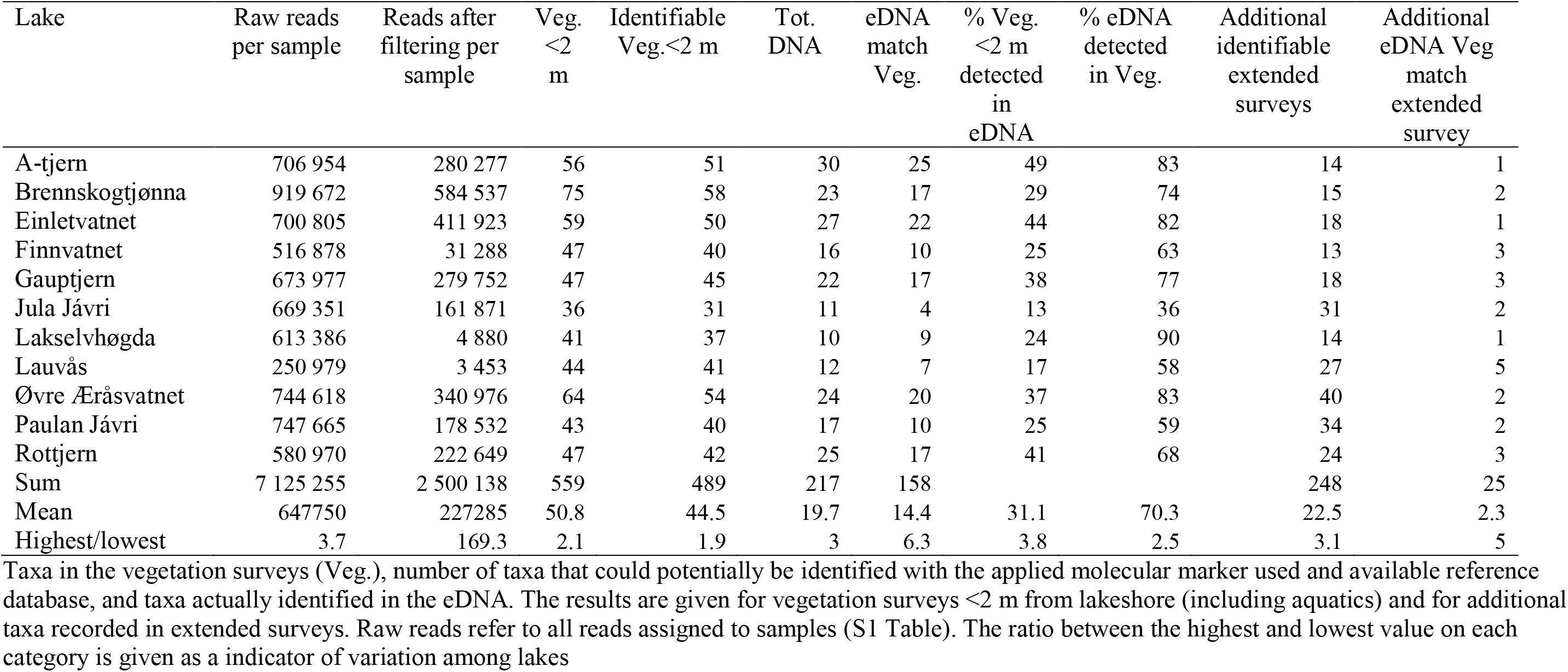
Number of records in vegetation and eDNA per lake.

Of 489 records <2 m from the lakeshore, the majority were rare (148) or scattered (146) in the vegetation; fewer were common (131) or dominant (64). An additional 245 observations of 46 taxa came from >2 m from the lakeshore (156 rare, 68 scattered, 19 common and 2 dominant).

### Molecular data

The numbers of sequences matching entries in the regional arctic-boreal and EMBL-r117 databases were 227 and 573 at 98% identity, respectively. For sequences matching both databases, we retained the arctic-boreal identification; this resulted in 11,236,288 reads of 301 sequences matched at 100% similarity with at least 10 reads in total (S2 Table). There were 244 and 181 records of taxa that with certainty could be defined as true or false positive, respectively (see methods). We found no combination of filtering criteria that only filtered out the false positives without any loss of true positives (S3 Table). The best ratio was obtained when retaining sequences that were on average more common in samples than in negative controls, plus with at least two replicates in one sample and at least 10 reads per replicate. Applying these criteria filtered out 163 false positives leaving only 18 records of three false positive taxa (Annonaceae, Meliaceae and Solanaceae), which were then removed as obvious contamination. However, it also removed 61 (25%) true positives, e.g., *Pinus*, which had high read numbers at lakes in pine forest and low ones at lakes where it is probably brought in as firewood, but which also occurred with high read numbers in two of the negative controls (S4 Table). After matching against the local vegetation, 2,500,138 reads of 56 sequences remained. Sequences representing the same taxa were merged, resulting in 47 final taxa (Table 3). Taking into account some taxa shared sequences, e.g. *Carex* and *Salix*, these may potentially represent 81 taxa (S1 Table).

**Table 3.**
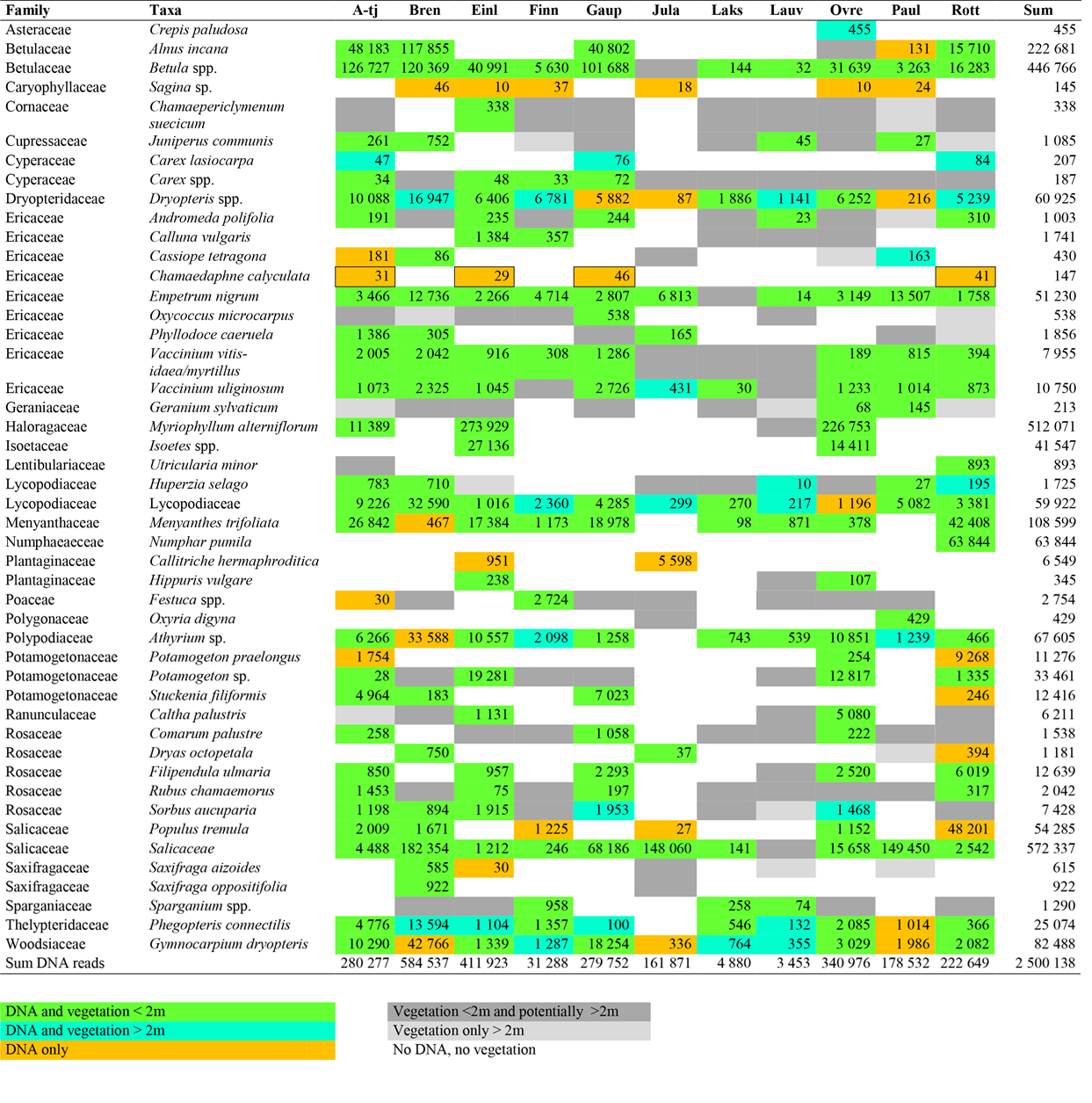
Read numbers per taxa and per lake.

The read numbers are sum of two DNA extractions with 6 PCR replicates for each. All read numbers are after the filtering steps in S2 Table. Note that the records of *Chamaeodaphne calyculata* are likely to represent false positives. For taxa only recorded in vegetation and/or filtered out of the eDNA records, see S1 Table. The lakes names are A-tjern (A-tj), Brennskogtjørna (Bren), Einletvatnet (Einl), Finnvatnet (Finn), Gauptjern (Gaup), Jula Jávri (Jula), Lakselvhøgda (Laks), Lauvås (Lauv), Øvre Æråsvatnet (Ovre), Paulan Jávri (Paul), and Rottjern (Rott).

In our positive control, 7 out of 8 species were detected in all replicates. Only *Aira praecox*, which was added with the lowest DNA concentration, could not be detected. This indicates that the PCR and sequencing was successful for taxa with an extracted DNA concentration of ≥0.03 ng/μL (S5 Table).

The gain in number of taxa when analysing two cores instead of one was 2.5±1.2 per lake. All data presented below are based on the upper 0-2 cm of sediment of two cores combined (but not from deeper levels as these were not sampled at all sites). This gave an average of 19.7±6.9 taxa (10-30) per lake (Table 2). Samples from below 2-cm depth provide an additional 14 records of 42 taxa, some not recorded in 0-2 cm samples (S1 Table).

### Detection of taxa in eDNA

Of the 217 eDNA records, the majority matched taxa recorded within 2 m of the lake shore (Figure 2a). Higher proportions of dominant or common taxa were detected in DNA compared with scattered or rare ones (Fig 2b). Most dominant taxa, such as *Betula, Empetrum nigrum, Vaccinium uliginosum*, and *Salix*, were correctly detected at most or all lakes (Table 3), whereas some were filtered out *(Equisetum* spp., *Pinus sylvestris*, many *Poa*, S1 Table). Of dominants, only two *Juncus* and two *Eriophorum* species were not recorded. Many taxa that were rare or scattered were filtered out (S1 and S4 Table).

**Fig 2.**
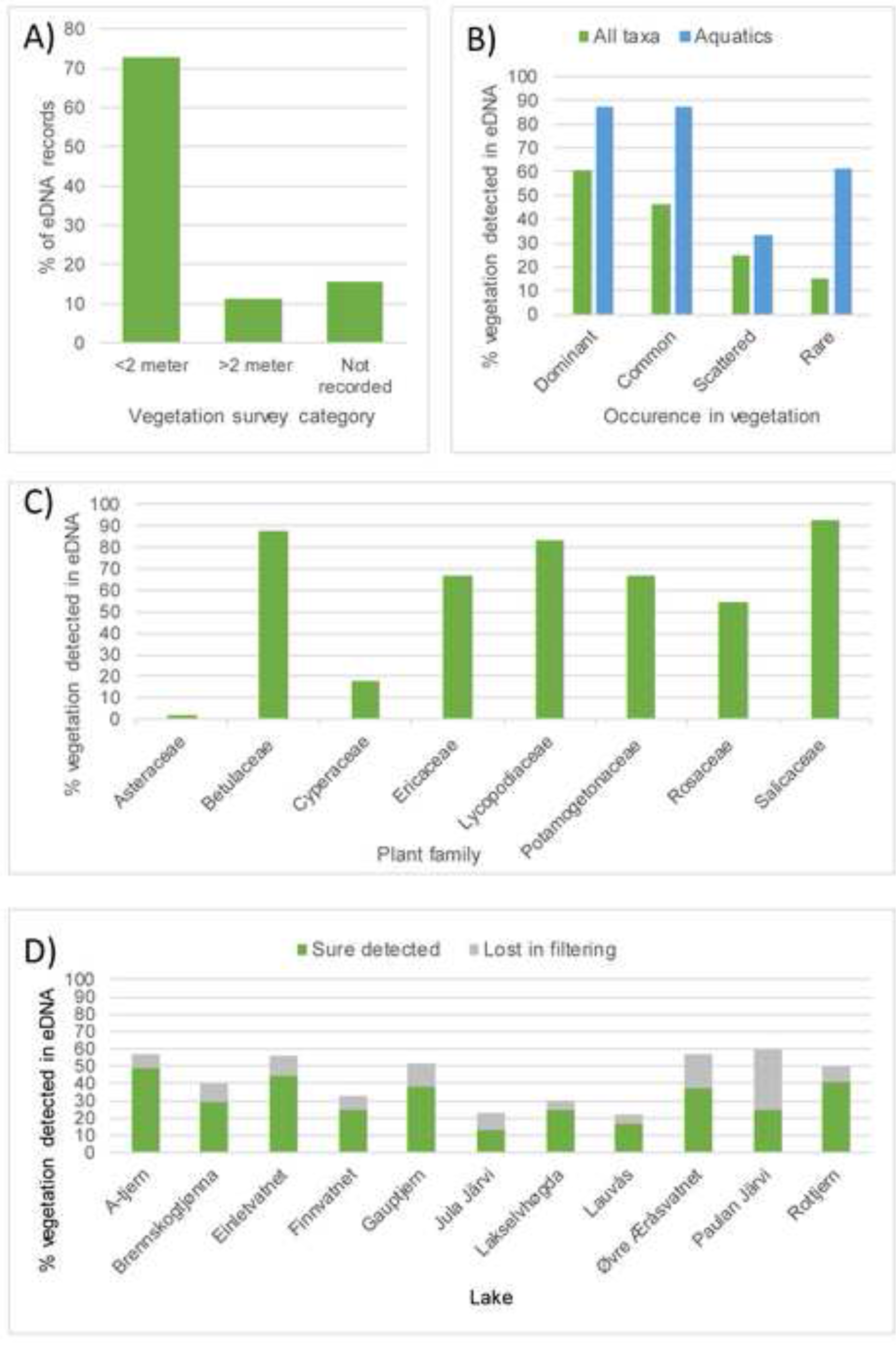
Match between records of taxa in the sedimentary eDNA in relation to vegetation surveys. a) Number of records in the sedimentary eDNA in relation to vegetation survey distance. b) Percentage records in eDNA in relation to abundance in vegetation surveys. c) Variation in percentage data among families with >11 eDNA records. d) Variation in percentage of taxa detected among lakes. Percentages in b), c) and d) refers to percentage of taxa recorded in the vegetation that potentially could be identified with the DNA barcode used. Note that DNA of more taxa were likely recorded but filtered out (S1-S4 Tables) – these numbers are only shown in figure b).

Detection success and taxonomic resolution in the eDNA varied among families (Table 3, Fig 2c). High success and resolution characterise Ericaceae and Rosaceae as they were identified to species level and successfully detected at most sites. Ferns (Dryopteridaceae, Thelypteridaceae, Woodsiaceae) and club mosses (Lycopodiaceae) were almost always detected, even when only growing >2 m from the lake shore. Aquatics (Haloragaceae, Lentibulariaceae, Menyanthaceae, Numphaeaceae, Plantaginaceae, Potamogetonaceae, Sparganiaceae) were also well detected, often also when not recorded in the vegetation surveys. Deciduous trees and shrubs (Betulaceae, Salicaceae) were also correctly identified at most lakes although often at genus level. In contrast, Poaceae and Cyperaceae, which were common to dominant around most lakes, were underrepresented in the DNA records. Juncaceae and Asteraceae, which were present at all lakes, although mainly scattered or rare, were mainly filtered out due to presence in only one PCR repeat or only in samples from 2-8 cm depth (S1-S4 Tables).

The numbers of taxa recorded in vegetation, in eDNA, and as match between them varied two- to six-fold among lakes (Table 2, Fig 2d). Jula Jávri had the lowest match between eDNA and vegetation with only four taxa in common. Lakselvhøgda and Lauvås had extremely low read numbers after filtering. For Lauvås, Finnvatnet and Lakselvhøgda, 84%, 30% and 20%, respectively, of raw reads were allocated to algae. If we assume that a big unidentified sequence cluster also represents algae, this increases to 69% for Lakselvhøgda, where a 15-20 cm algal layer was observed across most of the lake bottom. A lake-bottom algal layer was also observed at Jula Jávri, and in this we suspect that an unidentified cluster of 170,772 reads was algae. In most other lakes, algal reads were 3-15% (0.2% in Brennskogtjern, the lake with highest numbers of reads after filtering; algal data not shown).

Thirty-three records of 17 DNA taxa did not match vegetation taxa at a given lake (Table 3). These include taxa that are easily overlooked in vegetation surveys due to minute size (e.g., *Sagina* sp.), or only growing in deeper parts of the lake (e.g., *Potamogeton praelongus*). Other taxa are probably confined to ridge-tops of larger catchments, which lay outside the survey areas (e.g., *Cassiope tetragona* and *Dryas octopetala).* Two tree species that occur as shrubs or dwarf shrubs at their altitudinal limits, *Alnus incana* and *Populus tremula*, were found in the DNA at high-elevation sites. Also, ferns were detected at several sites where they were not observed in the vegetation surveys. On balance, most mismatches probably relate to plants being overlooked in the vegetation surveys or growing outside the survey area, whereas *Chamaedaphne calyculata* likely represents a false positive (Table 3, S1 Appendix).

The multivariate ordinations gave similar results for the vegetation and eDNA records with the only lake from Pine forest, Brennskogtjønna, and one of the two alpine lakes, Paulan Jávri, clearly distinguished on the first axis, whereas the lakes with varying cover of birch forest were in one cluster (Fig 3a-b). The other alpine lake, Jula Jávri, was only distinguished on the vegetation, probably due to the low number of taxa identified in the eDNA of this lake (Table 2). Percentages of variation explained by the first two axes were similar for the two analyses (CA Vegetation: Axis 1, □=0.50, 20.4%, Axis 2, □=0.37, 15.1%; eDNA: Axis 1, □=0.24, 18.9%, Axis 2, □=0.24, 18.5%). The Procrustes analyses indicated a good similarity between vegetation and eDNA (CA Correlation = 0.53, P=0.099; NSCA Correlation=0.59, P=0.045).

**Fig 3.**
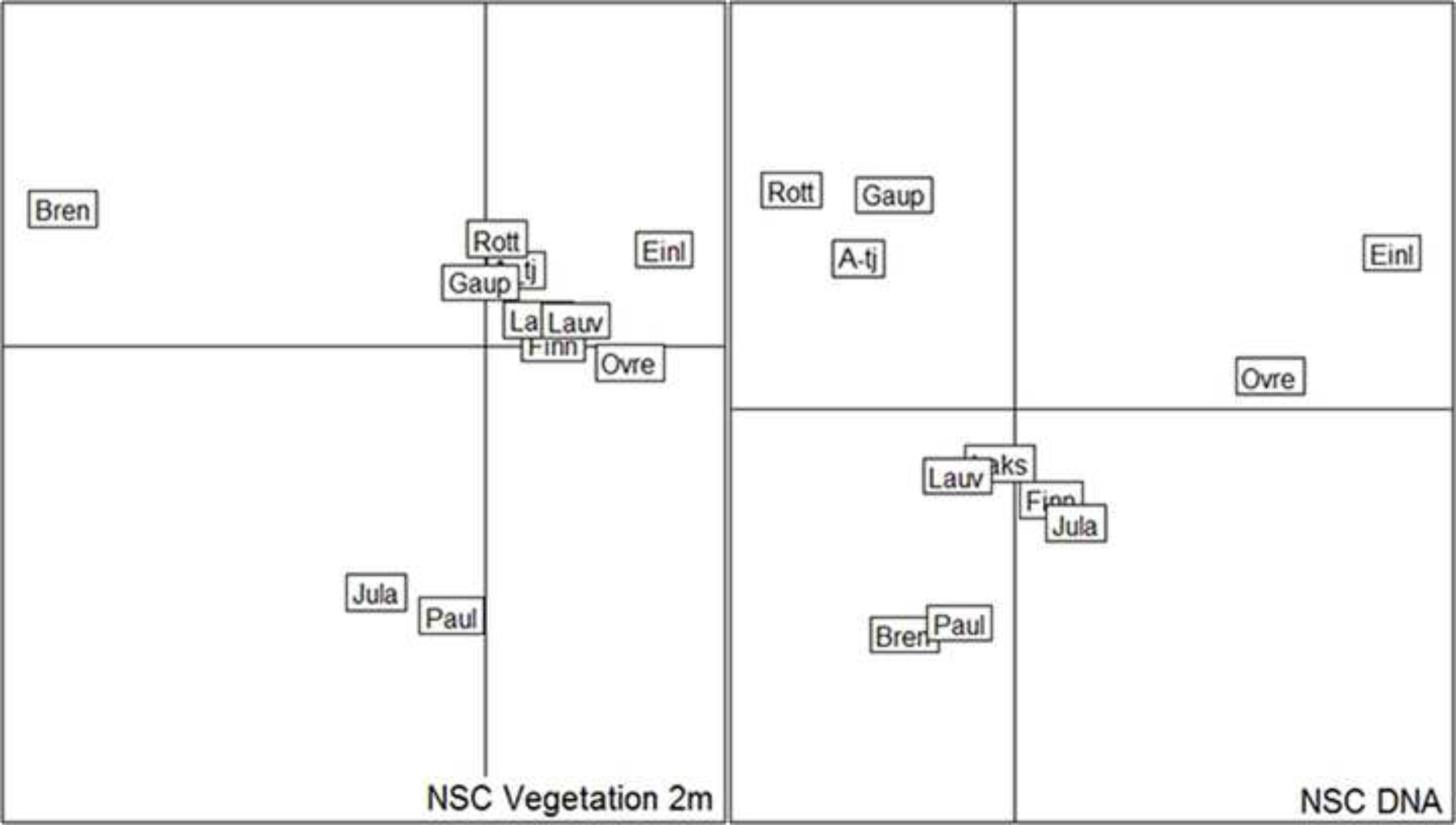
Multivariate ordination (Non Symmetric Correspondence Analysis; NSC) of the 11 lakes. The ordination is based on taxa recorded in the vegetation (a) and eDNA (b). Note that lakes in tall forbs birch/pine mixed forest (A-tjern, Rottjern, Gauptjern are clustered together in both plots; so are also Einletvatnet and Øvre Æråsvatnet (both mire/birch forest at the island Andøya), whereas some lake with poorer DNA records show some differences in clustering.

### Probability of detecting taxa in vegetation and DNA records

The posterior probability that all local taxa were recorded during the vegetation survey varied from 0.85-0.95 (S6 Table). Thus, on average, about three species may have been overlooked at each lake. The posterior probability that taxa recorded in the vegetation surveys and detected at least once by eDNA were also recorded in the DNA in a given lake (true positives) was 0.33-0.90, whereas the posterior probability of any DNA records representing a false positive varied from 0.06-0.33 per lake (S6 Table). There was evidence that the probability of detecting a species using eDNA (*p_DNA_1_*) was higher for deeper lakes (slope b_1_ = 0.58, 95% CI = [0.20; 0.98], Fig 4). Not surprisingly a similar effect was found for lake size (slope b_1_=0.25 [0.10, 0.41]) as lake size and depth were highly correlated (r=0.81). Catchment area (b_1_=0.06 [-0.15, 0.27]) and mean annual temperature (b_1_= - 0.03 [-0.14, 0.08]) did not appear to influence probability of detection by eDNA.

**Fig 4.**
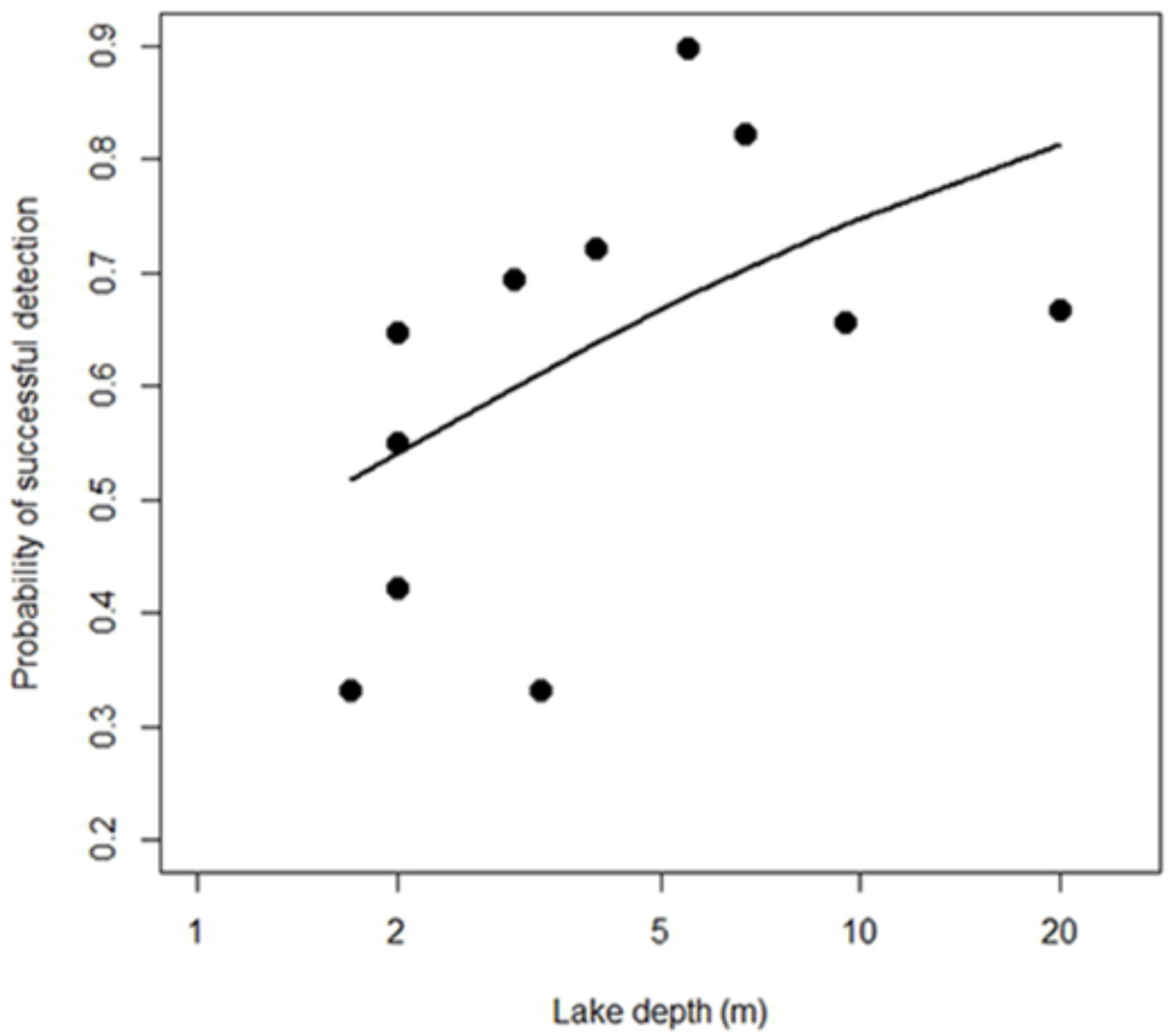
Lake depth versus detection probability. Relationship between lake depth and probability that a species present in the vegetation and detected at least once by eDNA is detected by eDNA in a given lake. The relationship is modelled as a logit function and back-transformed to the probability scale.

## Discussion

Taking into account the limitation of taxonomic resolution due to sequence sharing or taxa missing in the reference library, we were able to detect about one third of the taxa growing in the immediate vicinity of the lake using only two small sediment samples from the lake centre. The large number of true positives lost (S1 Appendix) suggests that this proportion may be further improved. Nevertheless, the current approach was sufficient to distinguish the main vegetation types.

### Taphonomy of environmental plant DNA

The high proportion of taxa in the <2 m survey detected with eDNA than in the extended surveys indicates that eDNA is mainly locally deposited. The observation of taxa not recorded in the vegetation surveys but common in the region (Fig 3, S1 Table) indicates that some DNA does originate from some hundreds of meters or even a few km distant. Indeed, a higher correlation between catchment relief and total eDNA (R^2^=0.42) than eDNA matching records in the vegetation (R^2^=0.34), may suggest that runoff water from snow melt or material blown in also contributes. Thus, the taphonomy of eDNA may be similar to that of macrofossils [66, 67], except that eDNA may also be transported via non-biological particles (e.g. fine mineral grains). From other studies, pollen does not appear to contribute much to local eDNA records [15, 35, 37, 42, 47]. This is probably due to its generally low biomass compared with stems, roots and leaves, and to the resilience of the sporopollenin coat, which requires a separate lysis step in extraction of DNA [68].

The higher proportion of eDNA taxa that matched common or dominant taxa in the vegetation, compared with taxa that were rare or scattered, was as expected, as higher biomass should be related to a greater chance for deposition and preservation in the lake sediments [9]. Yoccoz *et al*. [6] found the same in their comparison of soil eDNA with standing vegetation. While some dominant taxa were filtered out in our study, their DNA was mainly present (S1 Appendix, S1-4 Tables), and most dominant taxa were recorded in all PCR replicates (not shown). Thus, for studies where the focus is on detecting dominant taxa, running costs may be reduced by performing fewer PCR replicates.

### Variation among lakes

The variation among lakes seen in DNA-based detection of taxa shows that even when identical laboratory procedures are followed, the ability to detect taxa can vary. Our sample size of 11 lakes does not allow a full evaluation of the reasons for this variation. Factors such as low pH or higher temperature may increase DNA degradation [16], but the two lakes with lowest numbers of reads after filtering in our study, Lakselvhøgda and Lauvås, had pH values close to optimal for DNA preservation (7.2 and 6.8, respectively, I.G. Alsos and A.G. Brown, pers. obs. 2016), and variation in temperature was low among our sites. The lack of an inflowing stream at Lakselvhøgda may reduce the supply of eDNA, but Lauvås has two inflows. For these two lakes, and to a lesser extent Finnvatnet, we suspect high algal abundance might have caused PCR competition [69]. PCR competition may also occurred in samples from Jula Jávri, but in this case we were not able to identify the most dominant cluster of sequences. These lakes are also small and shallow. Variation among eDNA qualities has also been observed in a study of 31 lakes on Taymyr Peninsula in Siberia [70]. We suspect that high algae production may be a limiting factor as we also have seen poor aDNA results in samples with high Loss on Ignition values, but this should be studied further. A potential solution to avoid solution to avoid PCR competition may be to design a primer to block amplification of algae as has been done for human DNA in studies of mammals eDNA [71].

### Variation among taxa

The variation we observed among plant families, both in taxonomic resolution and likelihood of detection, is a general problem when using generic primers [45, 72, 73]. For example, the poor detection of the Cyperaceae may be due to the long sequence length of *Carex* and *Eriophorum* (>80 bp), and most studies only detect it at genus or family level [38, 42, 74]. The low representation of Asteraceae may be due to its rare or scattered representation in the vegetation and/or its poor amplification. While some studies successfully amplify Asteraceae [15, 37, 38, 42, 75], others do not, even when other proxies indicate its presence in the environment [46]. This may be due to the high percentage of Asteraceae taxa that have a one base-pair mismatch in the reverse primer [34]. Poaceae, which has no primer mismatch, is regularly detected in ancient DNA studies [15, 36-38, 41], and was present in nine lakes, although most records were filtered out due to occurrence in negative controls. To avoid any bias due to primer match and potentially increase the overall detection of taxa, one solution would be to use family-specific primers, such as ITS primers developed for Cyperaceae, Poaceae, and Asteraceae [36]. Alternatively, shotgun sequencing could be tested as this minimizes PCR biases [76, 77].

The common woody deciduous taxa *Betula* and *Salix*, as well as most common dwarf shrubs such as *Andromeda polifolia, Empetrum nigrum*, and *Vaccinium uliginosum*, were correctly detected in most cases. They are also regularly recorded in late-Quaternary lake-sediment samples [15, 25, 37, 41, 70, 74]. These are ecologically important taxa in many northern ecosystems, and their reliable detection in eDNA could be expected to extend to other types of samples, e.g., samples relating to herbivore diet [44].

The general over-representation of spore plants in eDNA among taxa only found >2 m from the lake and those not recorded in the catchment vegetation raises the question as to whether eDNA can originate from spores. Spore-plant DNA is well represented in some studies [42, 78], is lacking in other studies [15, 37] and has been found as an exotic in one study [41]. As with pollen, the protective coat and low biomass of spores suggest that they are an unlikely source of the eDNA. This inference is supported by clear stratigraphic patterns shown by fern DNA in two lake records from Scotland. Records are ecologically consistent with other changes in vegetation, whereas spores at the same sites show no clear stratigraphy [42]. Preferential amplification could be an alternative explanation, but this is not likely as the amplification of fern DNA from herbarium specimens is poor [34]. It is possible that in some cases, including this study, we are detecting the minute but numerous gametophytes present in soil, which would not be visible in vegetation surveys.

Aquatic taxa were detected in all lakes, and they have been regularly identified in eDNA analyses of recent [42] and late-Quaternary lake sediments [15, 37, 38]. eDNA may be superior to vegetation surveys in some cases, e.g., *Potamogeton praelongus*, which is characteristic of deeper water https://www.brc.ac.uk/plantatlas/) and was likely overlooked in surveys due to poor visibility. *Callitriche hermaphroditica* was observed in two lakes (Einleten and Jula javri), whereas *C. palustris* was observed at Einleten. We cross-checked the herbarium voucher and the DNA sequence and both seems correct, so potentially both were present but detected only in either eDNA or vegetation surveys. Overall, eDNA appears to detect aquatic plants more efficiently than terrestrial plants, which is not unexpected as the path from plant to sediment is short.

### The use of eDNA for reconstruction of present and past plant richness

In contrast to water samples, from which eDNA has been shown to represent up to 100 % of fish and amphibian taxa living in a lake [7, 79], one or two small, surficial sediment samples do not yield enough DNA to capture the full richness of vascular plants growing around a lake; the same limitation may apply in attempts to capture Holocene mammalian richness [22]. This is likely largely due to taphonomic limitations affecting preservation and transport on land, as aquatics were generally well detected. Also, surface samples are typically flocculent and represent a short time span, e.g. a few centimetre may represent 10-25 years ([49]; pers. obs.). Increasing the amount of material analysed, the amount of time sampled (by combining the top several cm of sediment), and/or the number of surface samples may improve detection rates for species that are rare, have low biomass and/or grow at some distance from the lake. In this study we identified more taxa when we used two surface samples and/or material from deeper in the sediment cores. Nevertheless, taphonomic constraints may mean that DNA of some species rarely reaches the lake sediment. On the technical side, both improvements in laboratory techniques and in bioinformatics could increase detection of rare species. In this study, DNA of many of the rarer taxa was recorded but was filtered out. As the rarest species are also difficult to detect in vegetation surveys [59], combining conventional and DNA-based surveys may produce optimal estimates of biodiversity.

The potential taxonomic resolution (i.e., for eDNA taxa to be identified to species level) was similar or higher than that for macrofossils [80] or pollen [81, 82]. The potential taxonomic resolution of any of these methods depends on how well the local flora is represented in the available reference collection/library, site-specific characteristics, such as the complexity and type of the vegetation [34, 82], and the morphological or genetic variation displayed by different taxonomic groups. In our case, only 3% of the taxa found in the vegetation surveys were missing in the reference database which likely improved the resolution. To reach 100% resolution, a different genetic marker is needed to avoid the problem of sequence sharing. Using longer barcodes may improve resolution [45, 83] and may work for modern samples, bur for taxa with cpDNA sharing as e.g. *Salix*, nuclear regions should be explored. For ancient samples with highly degraded DNA, taxonomic resolution may potentially be increased by using a combining several markers, hybridization capture RAD probe techniques, or full-genome approach [77, 84-86].

The actual proportion of taxa in the vegetation detected in the eDNA records (average 28% and 18% for <2 m and extended surveys, respectively, not adjusting for taxonomic resolution) is similar to the results of various macrofossil [80, 87-89] and pollen studies [81, 82]. This contrasts with five previous studies of late-Quaternary sediments that compared aDNA with macrofossils and seven that did so with pollen; these showed rather poor richness in aDNA compared to other approaches (reviewed in [10]). We think a major explanation may be the quality and size of available reference collections/libraries, as the richness found in studies done prior to the publication of the boreal reference library (e.g. [15, 27, 35, 37]) was lower than in more recent studies, including this one [42, 70, 90, 91]. The variation in laboratory procedures, the number and size of samples processed and the number of replicates also affect the results [4, 82, 86]. Nevertheless, the correlation between eDNA and vegetation found in the Procrust analyses show that the current standard of the method is sufficient to detect major vegetation types.

## Conclusion

Our study supports previous conclusions that eDNA mainly detects vegetation from within a lake catchment area. Local biomass is important, as dominant and common taxa showed the highest probability of detection. For aquatic vegetation, eDNA may be comparable with, or even superior to, in-lake vegetation surveys. Lake-based eDNA detection is currently not good enough to monitor modern terrestrial plant biodiversity because too many rare species are overlooked. The method can, however, detect a similar percentage of the local flora as is possible with macrofossil or pollen analyses. As many true positives are lost in the filtering process, and as even higher taxonomic resolution could be obtained by adding genetic markers or doing full genome analysis, there is the potential to increase detection rates. Similarly, results will improve as we learn more about how physical conditions influence detection success among lakes, and how sampling strategies can be optimized.

## Acknowledgements

We thank Marie Kristine Føreid Merkel, Veronica Rystad, Premasany Kanapathippillai, Chris Ware, Sarah Lovibond, Martin Årseth-Hansen, Torbjørn Alm and Antony G. Brown for field assistance, Department for Medical Biology at UiT for use of laboratory for extraction, Frederic Boyer for raw data handling, Torbjørn Alm for help with identifying plant specimens, and H. John B. Birks for valuable comments on an earlier draft of this manuscript.

## Supporting Information

**S1 Appendix.**
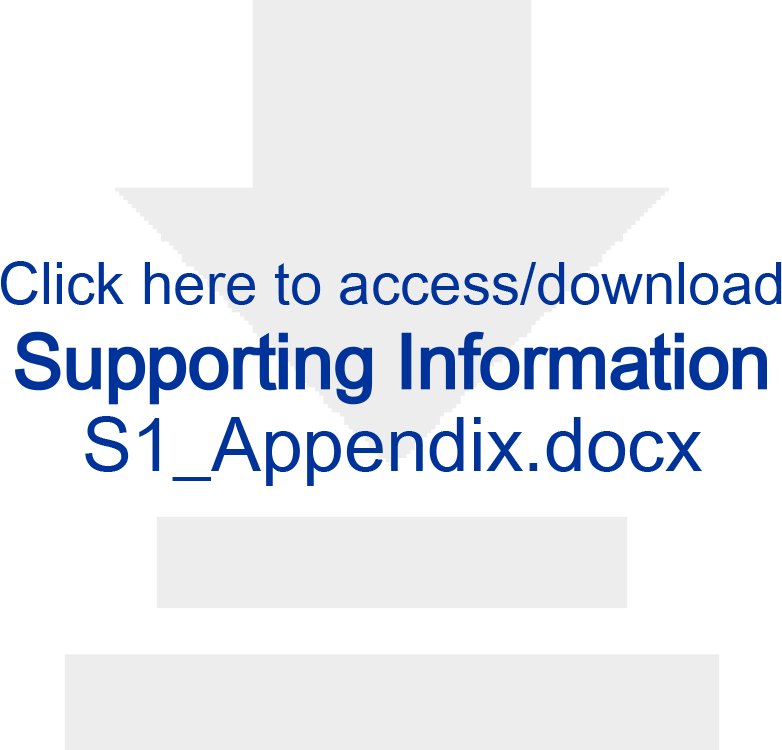
Comments on true and false positives.

**S1 Table.**
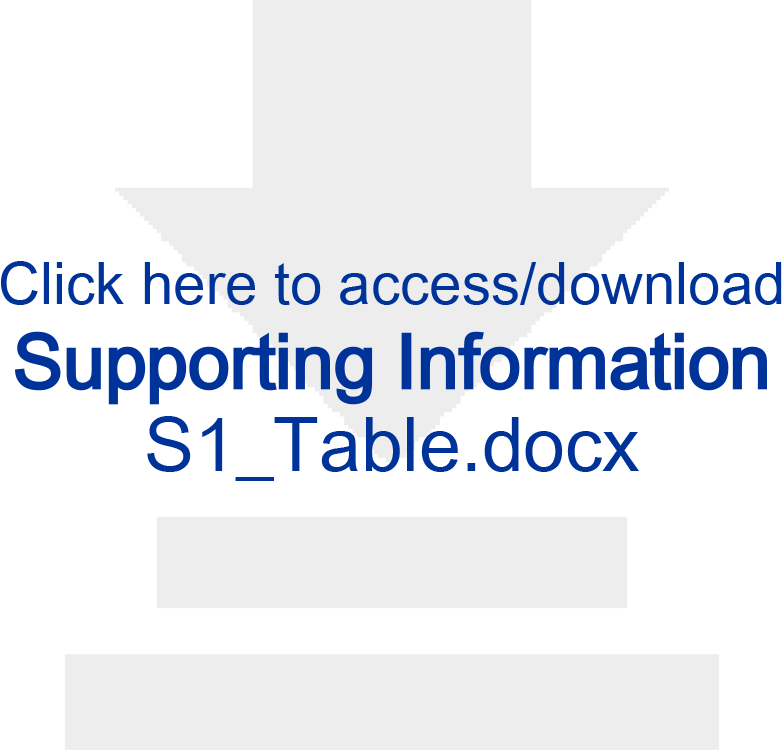
All taxa recorded in the vegetation surveys. (<2 m and/or larger surveys) at 11 lakes in northern Norway. Number refers to the highest abundance recorded among 2-17 vegetation polygons in the larger vegetation surveys (1=rare, 2=scattered, 3=frequent, and 4=dominant). Thus, 2316 records were combined to give one vegetation record per species and lake, in total 1000 records. Taxa match represent taxa that could potentially be identified by the molecular method used: ND=no data in reference library, ID incomp=could not be identified in DNA because the vegetation is incomplete identified, <12 bp= filtered out in initial filtering steps due to short sequence length. Max=the maximum abundance score observed at any of the lakes. The lakes names are A-tjern (A-tj), Brennskogtjørna (Bren), Einletvatnet (Einl), Finnvatnet (Finn), Gauptjern (Gaup), Jula Jávri (Jula), Lakselvhøgda (Laks), Lauvås (Lauv), Øvre Æråsvatnet (Ovre), Paulan Jávri (Paul), and Rottjern (Rott). See colour codes below. Hatched colour refer to DNA-vegetation match at higher taxonomic level (e.g. *Salix*).

**S2 Table.**
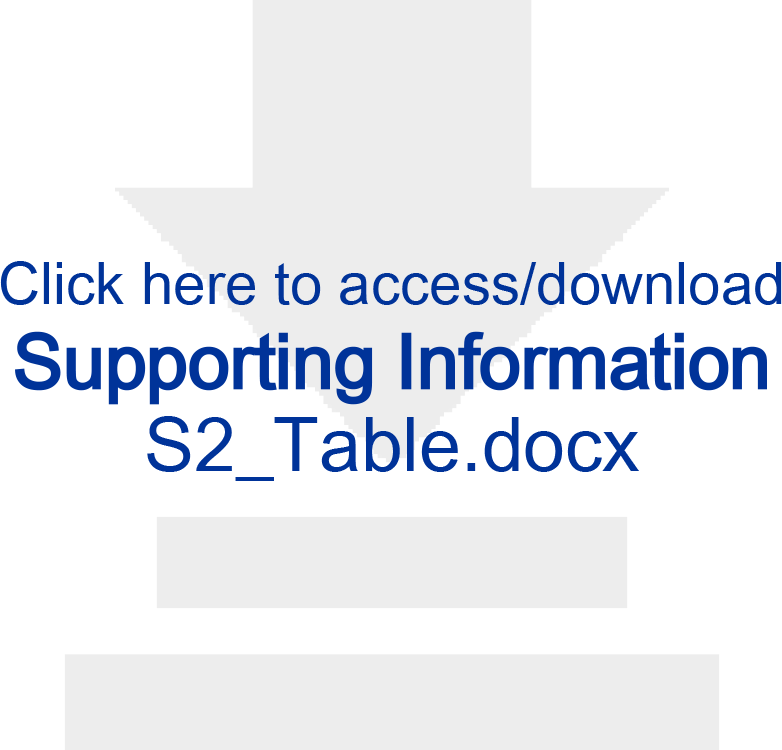
Number of sequence reads remaining after each filtering step for 42 samples from modern lake sediment collected in northern Norway, 6 extraction negative controls, 6 PCR negative controls and 2 PCR positive controls. Six individually tagged PCR repeats were run for each sample, giving a total of 336 PCR samples. Numbers of sequences and unique sequences are given for applying the criteria to all sequences.

**S3 Table.**
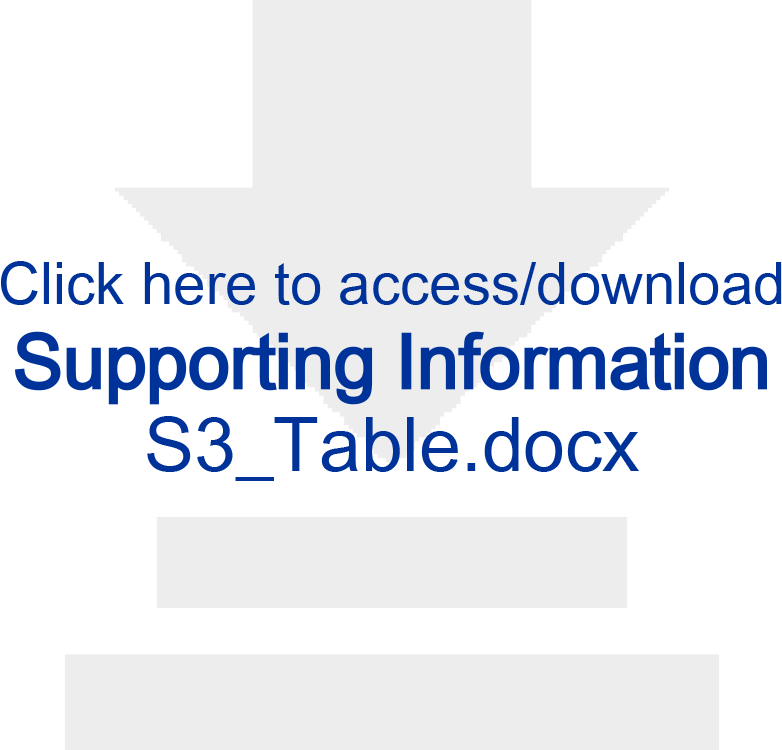
Effect of different filtering criteria on the number of True Positives. True positive (TP, defined as species also detected in vegetation surveys thus lower than the numbers given in Table 2) and False Positives (FP, defined as species not found in the regional flora; including 15 potential food plants) per lake and in total. The criteria used in this study, which gives the highest ratio between TP kept and FP lost, is shown in bold. 1) Minimum number of reads in lake samples, 2) minimum number of PCR repeats in lake samples, 3) “1” if occurring in more than 1 PCR repeat of negative control samples, 4) “1” if number of PCR repeats in lake sample > PCR repeats in negative control samples, 5) “1” if mean number of reads in lake samples > mean number of reads in negative control samples. The lakes names are A-tjern (A-tj), Brennskogtjørna (Bren), Einletvatnet (Einl), Finnvatnet (Finn), Gauptjern (Gaup), Jula Jávri (Jula), Lakselvhøgda (Laks), Lauvås (Lauv), Øvre Æråsvatnet (Ovre), Paulan Jávri (Paul), and Rottjern (Rott).

**S4 Table.**
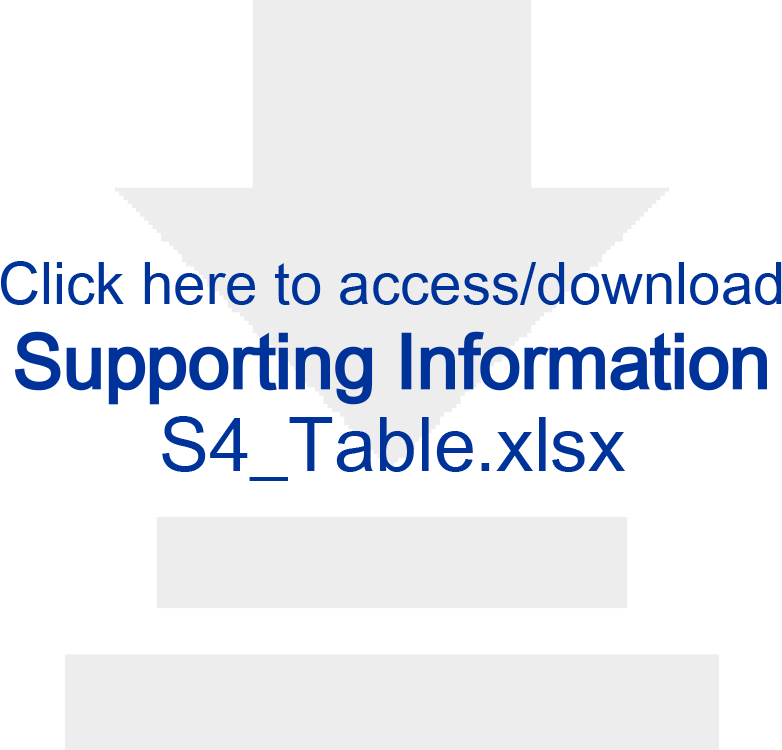
Taxa removed during filtering. All DNA reads that have 100% match to the reference libraries and have been removed during the second last step of filtering (see Table S2).

**S5 Table.**
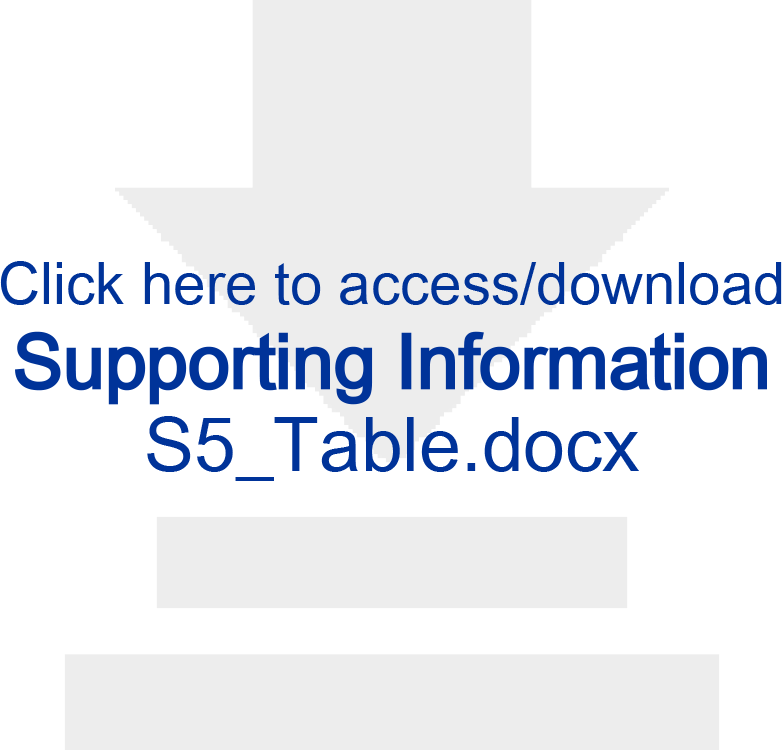
Retrieval of positive controls from raw Orbitool output file. The file consisted of 12706536 reads of 581 sequences, 98% match). Note that not all taxa used in the positive controls were present in the reference library but they match to closely related taxa.

**S6 Table.**
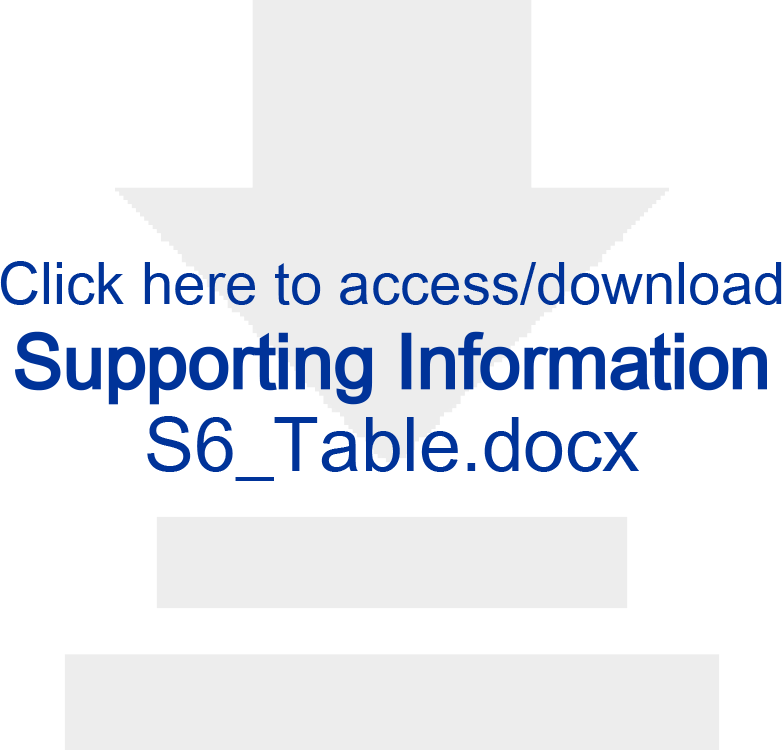
The probability of detection in eDNA and vegetation. The probability that all taxa in the vegetation were recorded (Vegetation), and that the DNA records represents true and false positives. Mean probability, standard deviation (SD) are given for each lake.

